# The eccDNA Replicon: A heritable, extra-nuclear vehicle that enables gene amplification and glyphosate resistance in *Amaranthus palmeri*

**DOI:** 10.1101/2020.04.09.032896

**Authors:** William T. Molin, Allison Yaguchi, Mark Blenner, Christopher A. Saski

**Author notes:** Corresponding Author: Christopher A. Saski. The author responsible for distribution of materials integral to the findings presented in this article in accordance with the policy described in the Instructions for Authors (www.plantcell.org) is: Christopher A. Saski.

## Abstract

Gene copy number variation is a predominant mechanism by which organisms respond to selective pressures in nature, which often results in unbalanced structural variations that perpetuate as adaptations to sustain life. However, the underlying mechanisms that give rise to gene proliferation are poorly understood. Here, we show a unique result of genomic plasticity in *Amaranthus palmeri*, a massive, ∼400kb extrachromosomal circular DNA (eccDNA), that harbors the 5-enoylpyruvylshikimate-3-phosphate synthase (*EPSPS*) gene and 58 other encoded genes whose functions traverse detoxification, replication, recombination, transposition, tethering, and transport. Gene expression analysis under glyphosate stress showed transcription of 41 of the 59 genes, with high expression of *EPSPS*, aminotransferase, zinc-finger, and several uncharacterized proteins. The genomic architecture of the eccDNA replicon is comprised of a complex arrangement of repeat sequences and mobile genetic elements interspersed among arrays of clustered palindromes that may be crucial for stability, DNA duplication and tethering, and/or a means of nuclear integration of the adjacent and intervening sequences. Comparative analysis of orthologous genes in grain amaranth (*Amaranthus hypochondriacus*) and water-hemp (*Amaranthus tuberculatus*) suggest higher order chromatin interactions contribute to the genomic origins of the *Amaranthus palmeri* eccDNA replicon structure.

**One-sentence summary:** The eccDNA replicon is a large extra-nuclear circular DNA that is composed of a sophisticated repetitive structure, harbors the *EPSPS* and several other genes that are transcribed during glyphosate stress.

## INTRODUCTION

McClintock (McClintock, 1984) stated “a sensing mechanism must be present in plants when experiencing unfavorable conditions to alert the cell to imminent danger and to set in motion the orderly sequence of events that will mitigate this danger.” Amplification of genes and gene clusters is an example of one such stress avoidance mechanism that leads to altered physiology and an intriguing example of genomic plasticity that imparts genetic diversity which can rapidly lead to instances of adaptation. This phenomenon is conserved across kingdoms, and these amplified genes are often incorporated and maintained as extrachromosomal circular DNAs (eccDNA). EccDNAs are found in human healthy and tumor cells (Kumar et al., 2017; Moller et al., 2018), cancer cell lines (Vanloon et al., 1994), in a variety of human diseases, and with limited reports in plants and other eukaryotic organisms (Lanciano et al., 2017). Recent research has reported that eccDNAs harbor bona fide oncogenes with massive expression levels driven by increased copy numbers (Wu et al., 2019). Furthermore, this study also demonstrated that eccDNAs in cancer contains highly accessible chromatin structure, when compared to chromosomal DNA, and enables ultra-long range chromatin contacts (Wu et al., 2019). Reported eccDNAs vary in size from a few hundred base pairs to megabases, which include double minutes (DM), and ring chromosomes (Storlazzi et al., 2010; Turner et al., 2017; Koo et al., 2018b; Koo et al., 2018a). The genesis and topological architecture of eccDNAs are not well understood, however, the formation of small eccDNAs (<2kb) are thought to be a result of intramolecular recombination between adjacent repeats (telomeric, centromeric, satellite, etc.) initiated by a double-strand break (Mukherjee and Storici, 2012) and propagated using the breakage-fusion-bridge cycle (McClintock, 1941). Larger eccDNAs, more than 100kb, such as those found in human tumors, most likely arise from a random mutational process involving all parts of the genome (Vonhoff et al., 1992). In yeast, some larger eccDNAs may have the capacity to self-replicate (Moller et al., 2015), and up to eighty percent of studied yeast eccDNAs contain consensus sequences for autonomous replication origins, which may partially explain their maintenance (Moller et al., 2015).

In plants, the establishment and persistence of weedy and invasive species has long been under investigation to understand the elements that contribute to their dominance (Thompson and Lumaret, 1992; Chandi et al., 2013). In the species *Amaranthus palmeri*, amplification of the gene encoding 5-enoylpyruvylshikimate-3-phosphate synthase (*EPSPS*) and its product, EPSP synthase, confers resistance to the herbicide glyphosate. The *EPSPS* gene may become amplified 40 to 100-fold in highly resistant populations (Gaines et al., 2010). Amplification of the *EPSPS* gene and gene product ameliorates the unbalanced or unregulated metabolic changes, such as shikimate accumulation and loss of aromatic amino acids associated with glyphosate activity in sensitive plants (Gaines et al., 2010; Sammons and Gaines, 2014). Previous low-resolution FISH analysis of glyphosate-resistant *A. palmeri*, shows the amplified *EPSPS* gene is distributed among many chromosomes, suggesting a transposon-based mechanism of mobility; while EPSP synthase activity was also elevated (Gaines et al., 2010). Partial sequencing of the genomic landscape flanking the *EPSPS* gene revealed a large contiguous sequence of 297-kb, that was termed the *EPSPS* cassette (Molin et al., 2017), and amplification of the *EPSPS* gene correlated with amplification of flanking genes and sequence. Flow cytometry revealed significant genome expansion (eg. 11% increase in genome size with ∼100 extra copies of the cassette), suggesting that the amplification unit was large (Molin et al., 2017). A follow up study reported that this unit was an intact eccDNA, and through high resolution fiber extension microscopy, various structural polymorphisms were identified that include circular, dimerized circular, and linear forms (Koo et al., 2018a). A comprehensive analysis of eccDNA distribution in meiotic pachytene chromosomes also revealed a chromosome tethering mechanism for inclusion in daughter cells during cell division, rather than complete genome integration. Here, we present the reference sequence of the eccDNA replicon, its unique genomic content and structural organization, experimental verification of autonomous replication, and a putative mechanism of genome persistence. We anticipate these findings will lead to a better understanding of adaptive evolution, and perhaps, toward the development of stable artificial plant chromosomes carrying agronomically useful traits.

## RESULTS

### Sequence composition and structure of the eccDNA plant replicon

Recent work determined the partial genomic sequence and cytological verification of the presence of a massive eccDNA that tethers to mitotic and meiotic chromosomes for transgenerational maintenance in glyphosate resistant *A. palmeri* (Molin et al., 2017; Koo et al., 2018a). Through single-molecule sequencing of the complete BAC tile path, our sequence assembly further corroborates the circular structure as a massive extrachromosomal DNA element (Koo et al., 2018a). The circular assembly is comprised of 399,435 bp, and is much larger than any eccDNA reported to date in plants (Figure 1A). The internal links connect direct and inverted sequence repeats with their match(es) indicating extensive repetitive sequences within the replicon (Figure 1A).

**Figure 1.**
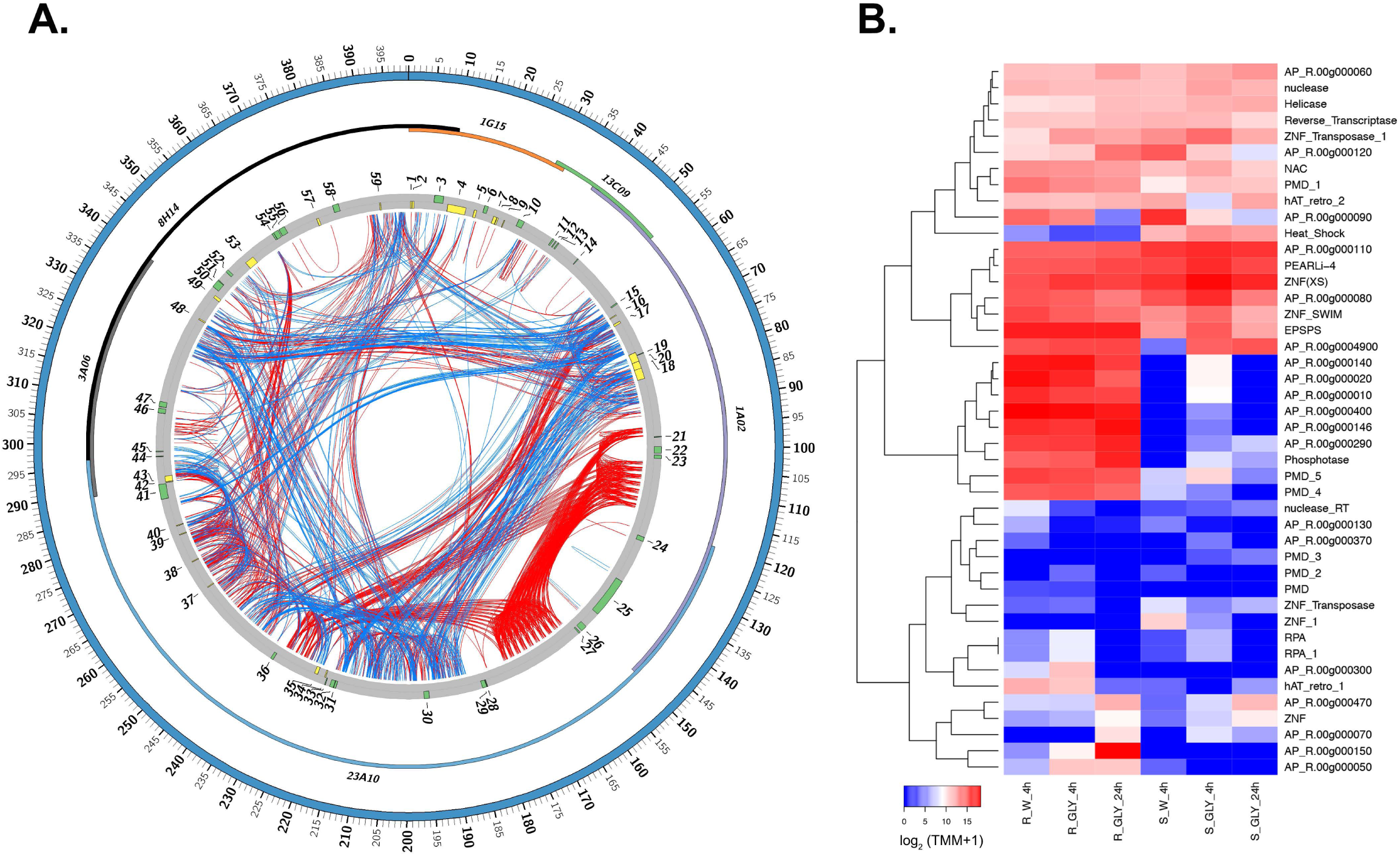
The eccDNA replicon structure and gene expression profiles. **A**. Circos plot of the *Amaranthus palmeri* eccDNA replicon. The solid blue ideogram represents the circular reference assembly of 399,435bp. The multicolored inner ideogram displays the overlapping alignment of sequenced BAC clones (1G15, 13C09, 1A02, 23A10, 3A06, and 8H14) composing the 399k assembly. The next track (gray) depicts the 59 predicted genes with their direction of transcription designated (green is clockwise, yellow is counterclockwise). The internal links connect direct (red) and inverted (blue) sequence repeats with their respective internal match(es). **B**. A heat map of gene expression of key genes under glyphosate (GLY) and water controls in a glyphosate resistant (R) and sensitive (S) biotype at 4 hours and 24 hours after treatment.

The eccDNA replicon contains 59 predicted protein coding gene sequences, including the *EPSPS* gene (Supplemental Table 1). The *EPSPS* gene is flanked by 2 putative AC transposase genes (Molin et al., 2017), which belong to a class of transcription factors known to interact with nucleic acid and other protein targets (Figure 1A, Supplemental Table 1). Nearby the *EPSPS* gene are two predicted replication protein A genes (Figure 1A, Supplemental Table 1), which can be involved in repair of double-strand DNA breaks induced by genotoxic stresses (Ishibashi et al., 2005). Downstream of *EPSPS* is a tandem array of genes with unknown functions but contain domains with Ribonuclease H-like activity and a C-terminal dimerization domain common to transposases belonging to the Activator (hAT) superfamily.

Many of the eccDNA replicon encoded genes have predicted functional protein domains that may endow the critical cellular processes necessary for stress avoidance, maintenance, stability, replication, and tethering of the eccDNA replicon. These processes include DNA transport and mobility, molecule sequestration, hormonal control, DNA replication and repair, heat shock, transcription regulation, DNA unwinding, and nuclease activity, (Supplemental Table 1). For example, the eccDNA replicon encodes 6 genes with aminotransferase domains (3 of which are highly expressed in the glyphosate resistant (GR biotype), which have been shown to be crucial for proper cell division and differentiation (Uhlken et al., 2014). The eccDNA replicon encodes a heat shock protein that belongs to the Hsp70 family, which is upregulated by heat stress and toxic chemicals; and may also inhibit apoptosis (Beere et al., 2000). There are 7 genes with plant mobile domains that function in the post-transcriptional gene silencing pathway, natural virus resistance, and the production of trans-acting siRNAs (Mourrain et al., 2000; Peragine et al., 2004) (RNAi machinery). A gene with a NAC domain is also predicted and expressed, which represents a class of transcription factors that regulate plant defense(Xie et al., 1999) and abiotic stress responses (Hegedus et al., 2003). There are 6 genes with zinc finger (ZF) domains, which are known to associate with DNA and protein, perhaps forming DNA-protein-DNA associations, that may function in chromosomal tethering. AP_R.00g000430 harbors a ZNF-SWIM domain, which has been shown to associate with *RAD*51 paralogs to promote homologous recombination (Makarova et al., 2002; Durrant et al., 2007). Furthermore, a recent study by Patterson et. al., 2019 has shown a RAD51 ortholog was co-duplicated with the *EPSPS* locus that confers glyphosate resistance in *Kochia scoparia* (Patterson et al., 2019); which may be indicative of parallel genomic recombination features. AP_R.00g000496 encodes a helicase domain, which is implicated in DNA replication and unwinding. AP_R.00g000450 contains a zinc binding reverse transcriptase domain and an integrase catalytic core, which are characteristic of a retroviral mechanism to integrate viral DNA into the host. This integrase is also found in various transposase proteins and is a member of the ribonuclease H-like superfamily involved in replication, homologous recombination, DNA repair, transposition, and RNA interference (Supplemental Table 1). In the GR biotype with an assembled eccDNA replicon, the most transcriptionally active genes at 24 hours post glyphosate treatment are a number of uncharacterized proteins, the *EPSPS* synthase (AP_R.00g000210), zinc finger nuclease (AP_R.00g000492), and several genes with plant mobile domains (Figure 1B).

### Transcriptional activity of the eccDNA replicon under glyphosate stress

Forty-one of the 59 genes encoded in the eccDNA replicon are transcriptionally active during at either 4 hours or 24 hours under glyphosate or water exposure in replicon containing glyphosate-resistant plants (Figure 1B and Supplemental Table 2). The most transcriptionally active genes include 2 uncharacterized proteins, followed by the *EPSPS* gene (Supplemental Table 2). Additional genes include 6 other uncharacterized protein sequences and 2 aminotransferase like genes that are mostly expressed in the GR biotype. Interestingly, at 4 hours after glyphosate and water control treatments, the resistant and sensitive plants gene expression profiles were indistinguishable from their water controls (Supplemental Figure 1 and Supplemental Table 2). At 24 hours after glyphosate exposure, 11 genes were detected to be upregulated in the GR biotype with at least a 2-fold increase when compared to the glyphosate sensitive biotype (Supplemental Table 3). These genes include 7 uncharacterized proteins, 2 aminotransferase-like genes, *EPSPS*, and a protein with a zinc finger SWIM domain (Supplemental Table 3).

### Repetitive elements and structural organization

The eccDNA replicon sequence is composed of a complex arrangement of direct and indirect repeat motifs of variable lengths dispersed among retroelements composed of SINEs, LINEs, and LTR elements, in addition to DNA transposons interspersed by predicted MITE and Helitron elements (Figure 2A and 2B). Flanking the *EPSPS* gene is an asymmetric set of direct repeats that are composed of arrays of clustered long and short interspersed palindromic repeat sequences (CLiSPrs), separated by identical MITES (Figure 2A). The complete palindromic array is bordered by LTR/ERVK, DNA/MULE MUDR and DNA/TdMar Stowaway elements (not shown). The CLiSPr block regions are also composed of elevated A+T segments (up to 80%), which may serve as a mechanism for stability or nuclear recognition sites for tethering, integration into open chromatin, or transcriptional hotspots. Downstream of the CLiSPr arrays are repetitive triplicate clusters of LTR-Cassandra and DNA hAT-Ac elements, each bordered by MITE elements. Still further downstream are clustered A-rich and LTR/Gypsy elements. These clustered arrangements may indicate functional relationships. The LTR/Cassandra and LTR/Gypsy elements divide the coding regions other than that contained within the direct repeats into 3 segments from 195 to 250 kb, from 287 to 355 kb and from 395 to 90 kb (Figure 3A and 3B).

**Figure 2.**
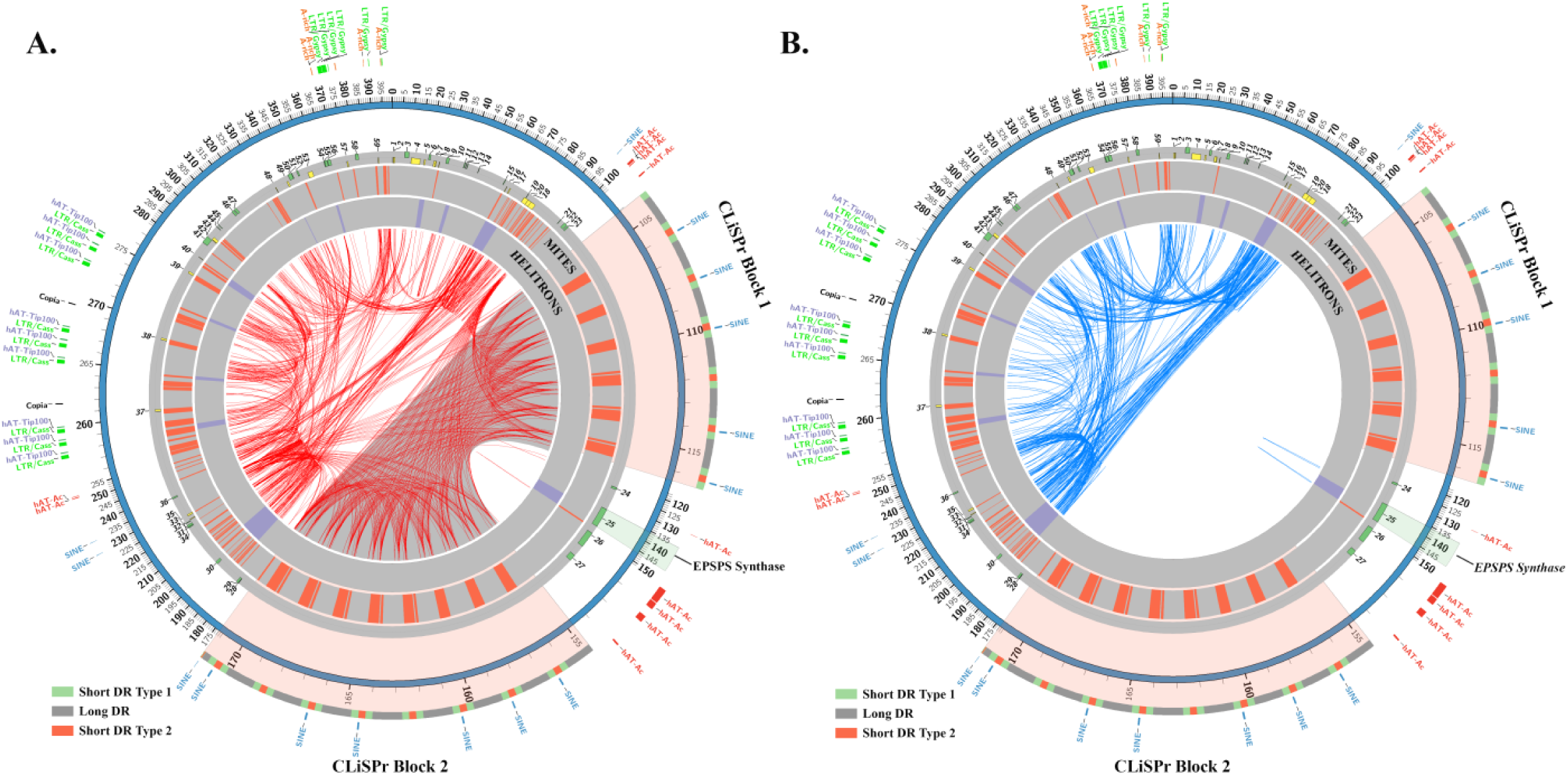
The eccDNA replicon repeat structure. **A**. Circos plot of the Amaranthus palmeri eccDNA replicon and key repeat content. The outer colored histograms with lables are predicted repetitive elements with organized arrangements. The highlighted repeat arrays are the clustered long and short interspersed palindromic repeats (CLiSPr) arrays that flank the EPSPS synthase gene The two repeat blocks are large and assymetric direct repeats. The innertracks depict predicted MITE and helitron repetitive elements. The internal red links are direct repeats and their relationships within the replicon. **B**. In this figure, the direct repeats are hidden and the indirect repeats are shown as blue links.

**Figure 3.**
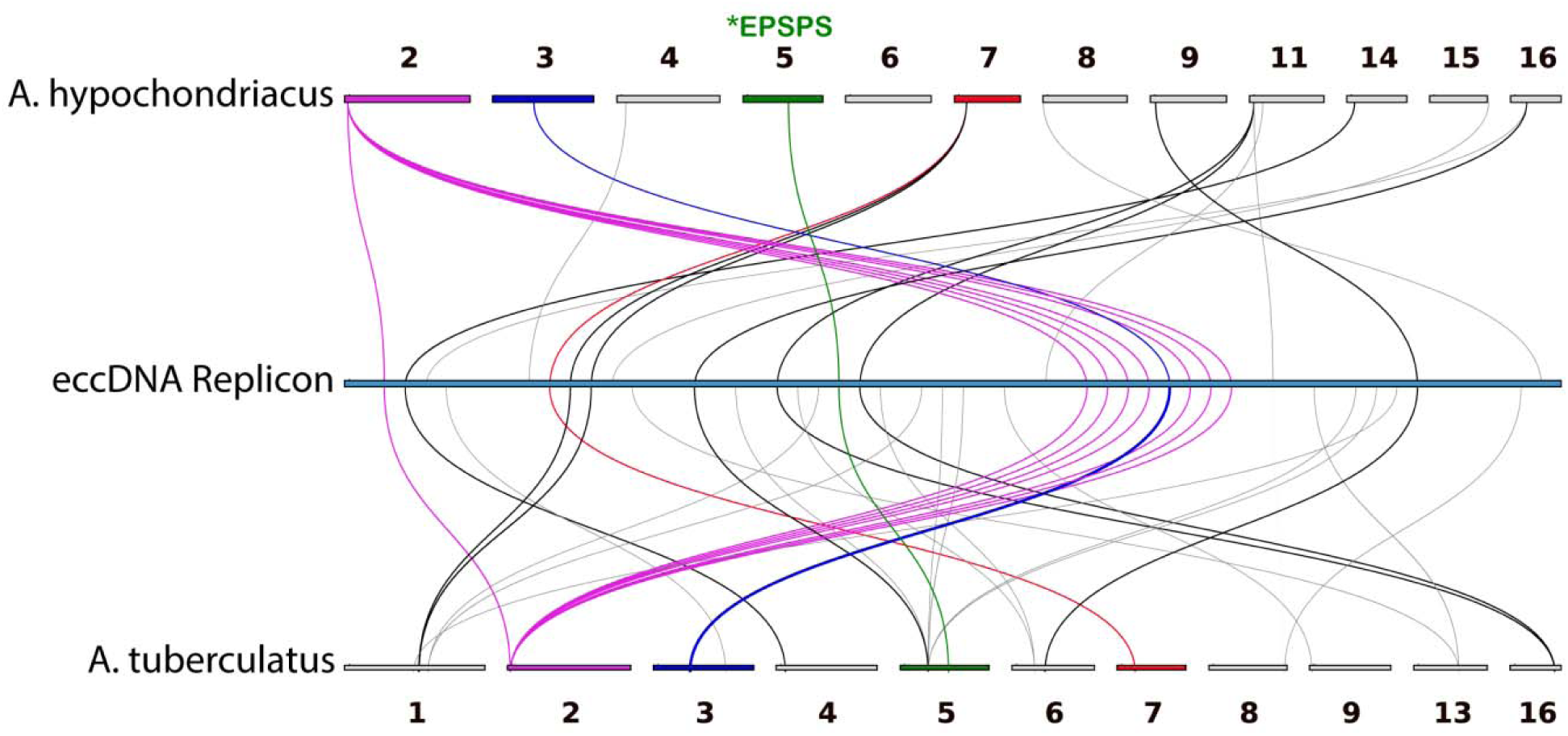
Collinear and syntenic arrangements in the A. hypochondracus and A. tuberculatus genomes and their relationship to the eccDNA replicon. Gray lines indicate syntenic gene arrangements between one or the other genomes and colored lines and ideograms indicate the gene is collinear in both genomes.

**Supplemental Figure 1.**
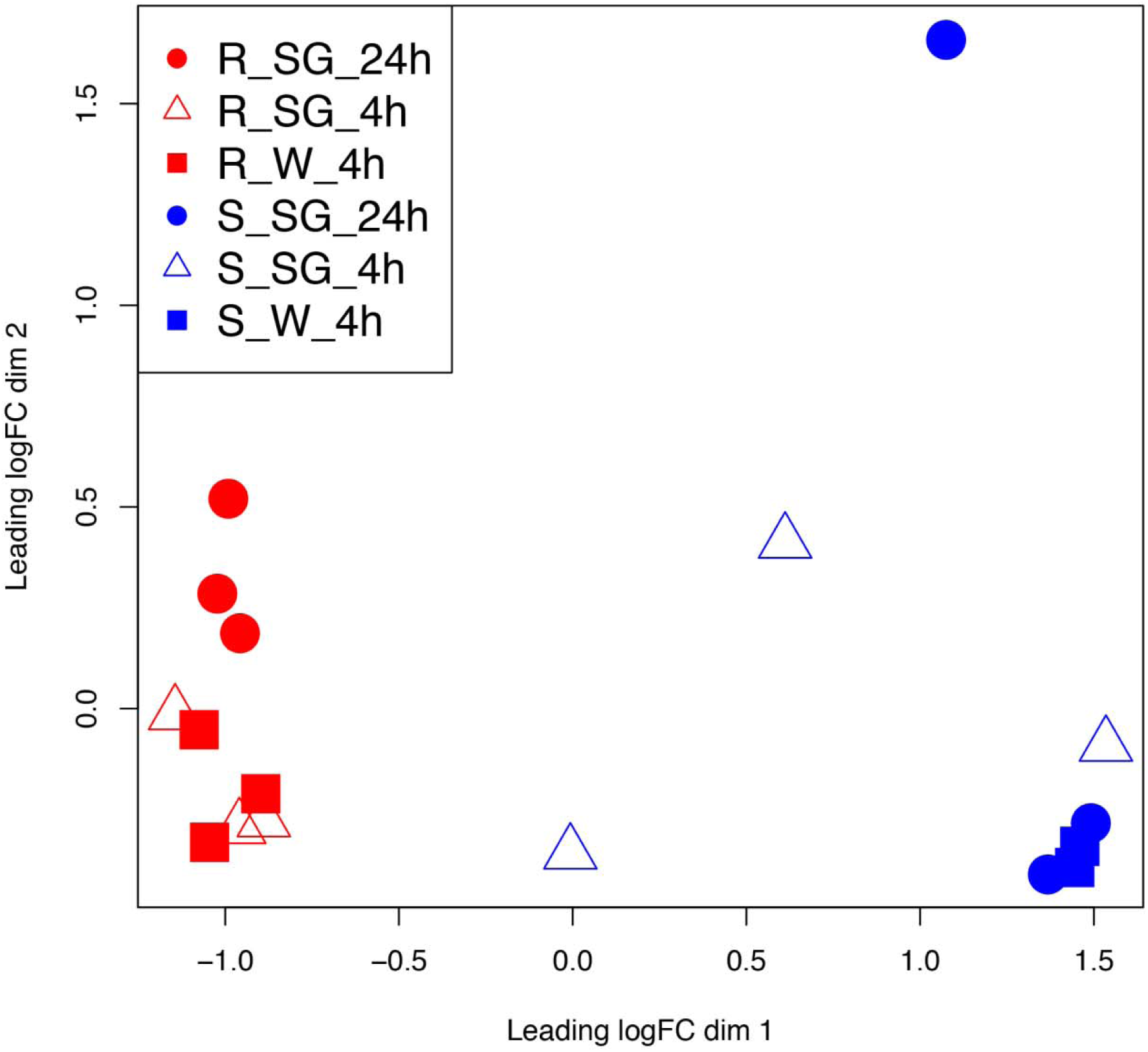
Multidimensional scaling plot of RNAseq data aligned to the eccDNA replicon reference derived from glyphosate resistant and sensitive biotypes sampled at 4 hours and 24 hours after exposure to glyphosate and water only controls. These data support Figure 1B.

### Genome tethering

The first cytological evidence of an eccDNA tethering to chromosomes in plants as a mechanism of genome persistence was recently reported (Koo et al., 2018b). EccDNA maintenance has been extensively studied in DNA viruses that maintain their genomes as extrachromosomal circular DNAs, such as Epstein-Barr, Rhadinovirus, papillomavirus, and others (Feeney and Parish, 2009). A commonality among these viruses is genome tethering, which is facilitated by virus encoded DNA-binding proteins that associate with repeated sequences in the viral genome that also have an affinity with host cell proteins that associate with mitotic chromatin to ensure nuclear retention (Feeney and Parish, 2009). For example, the Epstein-barr virus has been reported to anchor to the host genome through interaction of encoded Epstein-Barr nuclear antigen 1 (*EBNA1*) and the cellular protein Eukaryotic rRNA processing protein (*EBP2*) forming a protein-protein interaction or by directly associating to chromatin via an AT-hook motif that binds to A/T rich sequences on metaphase chromosomes (Wu et al., 2000; Sears et al., 2004). In Rhadinoviruses, a role has been suggested for the latency-associated nuclear antigen gene (*LANA*), which is thought to interact with terminal repeat regions (TR) in the virus (comprised of long tandem repeats) (Russo et al., 1996), where the C-terminus of the protein attaches the TR and the N-terminus tethers the episome to the chromosome (Piolot et al., 2001). In Papillomaviruses, the regulatory protein gene (*E2*) is a multifunctional DNA-binding protein that interacts with the helicase protein (E1 helicase) for replication (Masterson et al., 1998) and facilitates genome association through interaction within the N-terminal transactivation domain (Skiadopoulos and McBride, 1998). Computational analysis of the eccDNA replicon revealed several genes which may function in the tethering mechanism. AP_R.00g000496 contains 2 core AT-hook motifs (GRP) and also encodes a zinc finger SWIM domain which is recognized to bind DNA, proteins, and/or lipid structures (Supplemental Table 1). The optimal binding sequences of the core AT-hook are AAAT and AATT, which when bound together forms a concave DNA conformation for tight binding (Reeves, 2000). In the eccDNA replicion, there are 143 and 186 (AAAT)_2_ or (AATT)_2_ motifs, respectively (Figure 2A). There are consistent clusters of tandemly repeated motifs in the CLiSPr repeats. These AT-hook gene products may interact with the A/T rich regions of the eccDNA replicon and other nuclear scaffold proteins. Furthermore, the helicase domain has recently been demonstrated to be an important regulator of the chromatin association, establishment, and maintenance of the Herpes virus (Harris et al., 2017). AP_R.00g000496 is predicted to encode a helicase motif which may also have a role in tethering of the eccDNA to nuclear chromatin.

### Synteny and collinearity with Amaranthus hypochondriacus and Amaranthus tuberculatus

It is unknown whether the genesis of the eccDNA replicon was a result of a focal amplification surrounding the *EPSPS* gene or through recombination of distal genomic segments that combined multiple genes for simultaneous amplification. The high degree of synteny and collinearity preserved among the Angiosperms can offer insights into the genomic origin(s) of the eccDNA replicon. The eccDNA replicon was aligned to two the closely related species grain amaranth (*Amaranthus hypochondriacus*) and water-hemp (*Amaranthus tuberculatus*), each with a haploid chromosome number (n=16) and pseudochromosome-scale reference assemblies, (Lightfoot et al., 2017; Kreiner et al., 2019). Both comparator genome assemblies have a single copy of the *EPSPS* gene located near the middle of scaffold 5 (Figure 3). Out of the 59 eccDNA replicon genes, a total of 24 and 34 had reciprocal best hits (RBH) to the grain amaranth and water-hemp pseudochromosomes, respectively. The spatial topology of these ortholog matches was distributed across most of the pseudochromosomes in the assemblies (Figure 4 and Supplemental Table 4). Eleven of the RBH’s between the eccDNA replicon and the two *Amaranthus* genomes were located in a collinear fashion on the same scaffolds in the same order (Figure 4 colored ideograms and links). Interestingly, the first orthologous eccDNA gene (AP_R.00g000030 – aminotransferase-like) with synteny to scaffold 2 in both of the genomes is approximately 254 kb from the next cluster of collinear genes (Figure 4 – purple links) which are annotated in *A. hypochondriacus* as F-box family proteins with domain of unknown function (Supplemental Table 4). Moreover, this cluster of genes is separated in the middle by a gene that resides on a completely different pseudochromosome (scaffold 3) and is annotated as a NAC (No Apical Meristem) (Figure 4 blue link and Supplemental Table 4). These results suggest that the eccDNA replicon coding sequences likely originate from multiple genomic regions in *Amaranthus palmeri*, which supports the hypothesis of intra-genomic recombination during the formation of the eccDNA replicon.

The eccDNA replicon is a massive eccDNA vehicle for gene amplification, trait expression, maintenance and transfer of genomic information. The genomic origin is unknown, but likely a result of mobile element activation and extensive genome shuffling that may involve higher order chromatin interactions that may be influenced by xenobiotic pressures. It has various functional modalities for integration, stability and maintenance to ensure genomic persistence. Furthermore, because of the functional implications of the putative genes in the eccDNA replicon, the presence of this unit could cause a general increase in abiotic stress resilience, or perhaps, an increased disposition to adapt. This vehicle may afford new directions in breeding and biotechnology through deeper understandings of its origin and function. Further investigations will be required to ascertain the essential components of eccDNA replicon proliferation and its compatibility in other species.

## Methods

### Plant Material

The glyphosate-resistant (GR) *A. palmeri* plant used herein was collected in 2013 by W. T. M. from a soybean field in Washington County, Mississippi that was exposed to extensive application of glyphosate during the decade prior. Sixteen plants were collected and seeds collected from each plant were kept separate. For initial resistance assessment, seeds from each plant were sown in pots containing potting mix (Metro-Mix 360, Sun Gro Horticulture, Bellevue, WA) and lightly covered with 2 mm of mix and subirrigated. Pots were transferred to a greenhouse at the Jamie Whitten Delta States Research Center of USDA-ARS in Stoneville, Mississippi set to 25/20°C ± 3°C day/night temperature and a 15-h photoperiod under natural sunlight conditions supplemented with high-pressure sodium lights providing 400 µmol m^−2^ s^−1^. At the 2 leaf stage the seedlings were sprayed with glyphosate at 0.84 kg·ai·ha^−1^ using an air-pressurized indoor spray chamber (DeVries Manufacturing Co., Hollandale, MN) equipped with a nozzle mounted with 8002E flat-fan tip (Spraying Systems Co., Wheaton, IL) delivering 190 L·ha^−1^ at 220 kPa.. All seedlings survived from a plant designated #13 and seedlings from this seed lot were used for all experiments. #13 had a copy number as determined by qPCR of 34 (Molin et al., 2018).

### BAC Isolation, Sequencing, and Analysis

BAC library construction, partial tile path isolation, sequencing and analysis were described previously (Molin et al., 2017). Two additional BAC clones, 08H14 and 01G15, were determined by chromosome walking by hybridization with overgo probes designed from unique distal sequence on the terminal ends of the *EPSPS* cassette (clones 03A06 and 13C09). These two BAC clones were harvested and sequenced using Pacific Biosciences RSII sequencing to a depth greater than 100X, as described in (Molin et al., 2017). Raw single molecule sequence was self-corrected using the CANU Celera assembler (Koren et al., 2017) with the corOutCoverage=1000 to increase the output of corrected sequences. BAC end sequences were determined using standard Sanger sequencing methods and aligned to the reference assemblies with Phrap and opened in Consed (Gordon et al., 1998) for editing. BAC overlaps were identified using CrossMatch (Gordon et al., 1998) and ends joined manually to form a circular structure. The consensus eccDNA replicon was annotated using a combination of the MakerP pipeline (Campbell et al., 2014) with RNAseq (below) used as evidence with final manual curation. Functional domain scans and homology based annotations were determined by BLAST, InterproScan, and HMM using the SwissProt, non-redundant, and PfamA databases, respectively. Repeat characterization and masking were conducted with the RepeatMasker software (http://www.repeatmasker.org). MITE and helitron sequences were predicted with the detectMITE (Ye et al., 2016) and HelitronScanner (Xiong et al., 2014) tools. Circular figures were prepared using the Circos plotting toolset (Krzywinski et al., 2009).

### RNAseq

Plants for RNAseq were grown as previously to the two-true leaf stage under previously described greenhouse conditions at which time they were transplanted into 8 cm × 8 cm × 7 cm pots containing the same potting mix. Thereafter, plants were watered as needed and fertilized once two weeks after transplanting with a water-soluble fertilizer (Miracle-Gro, Scotts Miracle-Gro Products, Inc., Marysville, OH). When seedlings reached the six-leaf stage they were sprayed with either water, or water plus surfactant, or water plus surfactant plus glyphosate using the spray chamber. The surfactant was 0.5 % v/v Tween 20 and glyphosate was applied at 0.42 kg·ai·ha^−1^ after neutralization with 0.1 N KOH solution. Leaves from the third and fourth nodes were harvested for RNA extraction at 0, 4 and 24 hours after treatment. Plants were held for two weeks post leaf harvest to verify survival following glyphosate treatment.

Total RNA was harvested at 4 and 24 hours in biological triplicates using the RNeasy plant mini kit (Qiagen). Purified RNA was verified for intactness on a Bioanalyzer 2100 (Agilent) and subject to stranded mRNA-seq using standard TruSeq procedures and sequenced to a target depth of at least 15M reads per sample. Raw sequence data was preprocessed for adapter and low-quality bases with the Trimmomatic tool (Krzywinski et al., 2009) and cleaned reads aligned to the eccDNA replicon consensus assembly with Bowtie2 v.2.3.4.1 (Langmead and Salzberg, 2012) and the following arguments: --no-mixed --no-discordant --gbar 1000 --end-to-end –k 200 -q -X 800. TMM and FPKM transcript quantification was determined with RSEM v1.3.0 (Li and Dewey, 2011).

### Synteny and collinearity

Coding sequences (cds) and annotations (gff3) files for *Amaranthus hypochondricaus* (v1.0) were downloaded from Phytozome (https://phytozome.jgi.doe.gov). CDS and annotations for *Amaranthus tuberculatus* (v2, id54057) were downloaded from from CoGe (https://genomevolution.org/coge/SearchResults.pl?s=amaranthus&p=genome). Reciprocal best hits (RBH) were determined using the Basic Local Alignment Search Tool (BLAST) (Altschul et al., 1990) considering high identity paralogous matches. The linear riparian plot (Figure 3) was created with custom modifications to the python version of MCscan (https://github.com/tanghaibao/jcvi/wiki/MCscan-(Python-version)).

### Accession Numbers

Sequence data from this article can be found in the EMBL/GenBank data under BioProject ID PRJNA413471; Submission MT025716.

## Author Contributions and Acknowledgements

### Author contributions

W.M. and C.S. designed, conducted, analyzed, and interpreted the eccDNA replicon capture, sequencing, and analysis components of the study, which include BAC screening, isolation, sequencing, assembly, annotation, and data analysis. W.M. and C.S. also prepared the initial draft of the manuscript and edited subsequent versions. A.Y. and M. B interpreted data and edited the manuscript.

## Supplemental Data

Supplemental Table 1. Predicted coding genes and functional annotation of the eccDNA replicon.

Supplemental Table 2. Transcriptional abundance (log2 TMM+1) of eccDNA replicon genes under water and glyphosate exposure 4 hour and 24 hours after treatment in a glyphosate sensitive and resistant biotype.

Supplemental Table 3. Genes upregulated in glyphosate resistant biotype compared to glyphosate sensitive biotype 24 hours after glyphosate exposure. Numerical values included are normalized counts.

Supplemental Table 4. Replicon gene ortholog matches and locations in *A. hypochondriacus* and *A. tuberculatus* pseudochromosome assemblies.

